# Citizen science suggests decreased diversity of insects in Mexico, a megadiverse country

**DOI:** 10.1101/2025.07.01.662623

**Authors:** Jorge Soberon

**Affiliations:** Biodiversity Institute, University of Kansas, Dyche Hall, 1345 Jayhawk Blvd, Lawrence, KS 66045USA

## Abstract

An analysis of iNaturalist data on several taxonomic groups of insects in Mexico is presented. I found evidence of a decreasing trend in diversity of species per year, for four families of butterflies, the bumblebees and the dragon and damselflies. I performed the anlayses on several of the Potential Vegetation types of J. Rzedowsky, and explore the role of deforestation and pesticide use on the trends I found. I discuss the challenges of using unsystematic data to estimate trends and provide several hypotheses to explain them.

## Introduction

There is evidence of a decrease in insect populations worldwide (Hallmann et al. 2017, Edwards et al. 2025). This is worrisome among other things because insects are key components of ecosystems and provide societies with important ecosystem services, like pollination (Potts et al. 2010). Moreover, insects have a large and mostly unappreciated cultural importance (Duffus et al. 2021). This is true for the entire world, but even more so in countries like Mexico, where insects have a culinary use (Ramos-Elorduy and Viejo Montesinos 2007), are depicted by ancestral cultures (Beutelspacher 1989) and have an increasing social value (Rogel-Fajardo et al. 2011).

Most of the detailed evidence of the decline comes from the countries in the temperate zones that developed formal monitoring schemes (Sánchez Herrera et al. 2024). Tropical regions are less well studied, and evidence is contradictory (Wagner et al. 2021, Bonadies et al. 2024, Boyle et al. 2025). For instance, (Lucas et al. 2016), working on Hemiptera, and (Basset et al. 2017) on saturniid moths report no trends in Barro Colorado island, Panama. In Veracruz, Mexico, well monitored fruit flies show no trends (Aluja et al. 2012, Ordano et al. 2013). On the other hand, decreases in saturniid moths larvae have been reported in Costa Rica (Salcido et al. 2020); and declines in arthropod biomass have been reported in Puerto Rico, and with a few data points in Chamela, Mexico (Lister and Garcia 2018). The Monarch butterfly, perhaps the best monitored insect species in the country, shows consistent decreases in its wintering aggregations (Vidal and Rendón-Salinas 2014, Thogmartin et al. 2017, Zylstra et al. 2021).

Due to its history, climate, topography and cultural milieu (Ramamoorthy et al. 1993), Mexico is one of the megadiverse countries of the world, (Mittermeier et al. 1997), therefore, assessing the trends of insect populations in Mexico should be a priority. Unfortunately, long-term insect monitoring in Mexico is exceptional. Although in the country the history of entomological research is long, there are many collections, and hundreds of publications (Michán and Llorente 2002), monitoring is limited to a handful of species. Discovering the reasons for a lack of national monitoring schemes, like those existing in other countries (Thomas 2005, Streitberger et al. 2024) is worth doing, although I do not attempt it here. For Mexico, I take as the premise that there is only a handful of available insect time series. By that I mean time series more than two or three years long and systematically obtained. Since there are no national monitoring efforts, is it possible then to estimate insect’s biodiversity trends in Mexico?

Systematic monitoring efforts are compiled in one of a few world-wide population numbers time-series databases, like the Living Planet Index (Almond et al. 2020), or the Global Population Dynamics database (NERC Centre for Population Biology 1999). Unfortunately, such databases are sparse in insect information and contain no data for Mexico. Another possibility is to use so-called citizen science data (Cohn 2008), which although opportunistic and unsystematic, tends to be abundant. Data --on insects--collected by non-professionals has been used to estimate phenology and distributions (Soroye et al. 2018). Using it to estimate population trends is more challenging, as I discuss below.

In Mexico, the best citizen’s science initiative is iNaturalist (or Naturalista, as it is known in Mexico). iNaturalist in Mexico began its operations in 2008, although in 2013 the initiative came under the leadership of the national biodiversity agency, CONABIO, under the name of Naturalista (Macías and Freire 2017) and obtained funding from the Slim Foundation. Therefore, in Mexico, in some sense iNaturalist began in earnest after 2013. Despite the relatively late start, just for four families of butterflies, the damselflies, dragonflies and bumblebees, iNaturalist for Mexico contains more than 80,000 records tagged as “research level.” This is a substantial amount of information that may be used to assess trends. However, citizen science data must be corrected for biases. Specifically, in iNaturalist there are more observers and observations every year, at least in Mexico (see tables S1 and S2), and this bias should be considered when using such data.

Indeed, one of the main problems of using opportunistic citizen science data to estimate trends, is to correct for biases in recording effort (Di Cecco et al. 2021). There are several ways of dealing with this problem (Isaac et al. 2014, Outhwaite 2019, Tang et al. 2021). One of the “simplest” is correcting bias by obtaining the quotient of the metric used to report abundance to some measure of the effort invested in a locality, for a period. What is “effort” and how can it be measured using iNaturalist data? Collecting effort is difficult to define. It can be done in terms of time spent collecting, or number of individuals collecting. The iNaturalist data allows extracting a measure of time (number of monthly observations in a year) but their quality, beyond the tag of “research”, which refers to the reliability of the name assigned to the species, remains unreported. Since it has been suggested that the number of observations is preferable to the time spent observing (Willott 2001), in this work I will use, as measure of effort, the number of observers that have two or more observations. In each case the quality of the methodologies used, and the skill of the observers ideally should be known. Unfortunately, in the case of iNaturalist such quality is known to change (Di Cecco et al. 2021). Di Cecco (2021) suggests that, in iNaturalist, observers with at least two observations are more reliable than those with just one. Therefore, as a measure of effort I use the number of observers with two or more observations. The question then is to test for a trend in the diversity index with time. The naïve way of doing this would be by regressing the diversity index against time.

Ordinary regressions of metrics against time often suffer of the problems of autocorrelated errors and non-equal variances (heteroskedasticity), that are characteristic of time series (Shumway and Stoffer 2005), and this should be accounted for. One way of dealing with the complexities of analyzing a time-series, when the data are counts, is to use a package like ‘trim’ (in the platform R), which assumes a Poisson model for the underlying data (Pannekoek J. 1998). This approach corrects for the autocorrelation of errors and also for heteroskedasticity. Trim has been used frequently with European data (van Strien et al. 2019), and in tropical America (Novoyny and Y.Basset 2000), but the key assumption that the data are counts (a discrete scale) makes it difficult to analyze continuous-scale indices, or data with many non-occurrences, because the software is sensitive to the presence of zeroes, or NAs in the data.

Another possibility is to estimate whether there is a significant trend in the data, using a non-parametric Mann-Kendall test (Lyubchich et al. 2013). One first performs an ordinary least squares linear regression (OLS) of metric against time and then checks for the existence of trends (linear or monotonous). This method uses the sign of the slope in the OLS and the significance is tested by the Mann-Kendall test.

It is also possible to use, for a single species, the logit of the probability of occupancy of a cell, on a time unit (van Strien et al. 2019) and use Generalized Linear Modelling (glm) to fit a linear combination of the predictors, using as a measure of effort the length of the list of species (Szabo et al. 2010). Then several single species regressions are combined in an index (van Strien et al. 2019). One problem with this approach is that Generalized Linear Modelling has as assumptions of homoscedasticity and independence of errors, both of which may be violated in time series.

A statistically more sophisticated modification of the above idea is reporting the proportion of occupied sites under a hierarchical model that separates actual presence from the act of observing it (Outhwaite 2019). Although apparently very rigorous, this approach has its own problems, among them the need to define an appropriate model for the “present” and “observer” components, the need to define a way of aggregating observations in some grid of cells for the region, and ways to estimate the parameters of the observation part of the hierarchy.

Finally, another possibility is to resort to generalized least squares regressions (GLS), that allow for autocorrelated errors and heteroskedasticity. The R package ‘nlme’ implements this technique. One can then: fit (1) an ordinary regression of the index against covariates like time, and (2) another regression that includes in its model autocorrelation (the simplest ARIMA model of lag=1) with power variance decay. Then one can compare the two models using an Akaike criterion, and keep the best model. This is the option I used.

I also wanted to assess whether any existing trends were different for different ecological regions of Mexico. There are a variety of subdivisions of Mexico, from different ecological perspectives, and at different spatial resolutions (Miranda and Hernández-X 1963, Anonymous 1997, Olson et al. 2001, Challenger and Soberón 2012) I decided to use Rzedowsky’s Potential Vegetation types Rzedowsky (1986), that although coarse-grained, is based mostly on straightforward floristic criteria, it is well known in Mexico, and has a small number of categories.

An important caveat when using citizen science data is that species that are difficult to identify by sight should be avoided. In this work we focus on (1) three families of butterflies (Papilionidae, Pieridae, and Nymphalidae, with 316 names. We do not use Skippers or the smallest families in the Papilionoidea); on 23 names of bumblebees, and on 293 names for the Odonata (both Zygoptera and Anysoptera). Also, as a comparison, we included data on 307 names of the Solanaceae. The number of names (without proper taxonomic validation by experts), as reported by iNaturalist, appear in table 1.

**Table 1.** Number of different scientific names for the different taxonomic groups on the four most visited potential vegetation classes. The butterflies are the Papilionidae (swallowtails), Pieridae (sulphurs) and the Nymphalidae (brushfoots).

|  | All<br>Mexico | Xerophitic<br>Shrub | Pine Oak<br>Forest | Grasslands | Tropical<br>Deciduous<br>Forest |
| --- | --- | --- | --- | --- | --- |
| Butterflies | 413 | 153 | 220 | 220 | 198 |
| Odonata | 292 | 185 | 187 | 105 | 184 |
| Bombus | 23 | 16 | 20 | 9 | 14 |
| Solanaceae | 307 | 176 | 210 | 111 | 153 |

**Table 1.** Digital Object Identifiers and number of iNaturalist (“research” tagged) records downloaded, from the GBIF site, for the different taxonomic groups.

| TAXON | GBIF DOI | iNaturalist records | Unique names |
| --- | --- | --- | --- |
| Nymphalidae | doi.org/10.15468/dl.qta4zp | 20,145 | 223 |
| Papilionidae | doi.org/10.15468/dl.uu4unc | 5,583 | 13 |
| Pieridae | doi.org/10.15468/dl.zug4ee | 29,994 | 80 |
| Lycaenidae | doi.org/10.15468/dl.xgad6g | 2,326 | 97 |
| Odonata | doi.org/10.15468/dl.atdkfz | 36,775 | 292 |
| Anisoptera |  | 24,003 | 163 |
| Zygoptera |  | 12,772 | 129 |
| Bombus | doi.org/10.15468/dl.c6h4jz | 6,543 | 23 |
| Solanaceae | doi.org/10.15468/dl.597nj5 | 39,014 | 307 |

For the butterflies, it would be an interesting idea to use the species identified as indicators of “conservation” status (Orta et al. 2022), but most of the species mentioned by these authors have just a handful of records in the iNaturalist database, therefore I decided to perform the analysis pooling the three families of butterflies.

To obtain uncertainty bands, I define grids of hexagons at several resolutions, covering the territory of Mexico (see figure S1). Thus, for a given year and taxonomic group, I took means and variances over hexagons. Each unique combination of year and hexagon defines an “event”, and thus my metrics of abundance will be: (1) the number of observations per event (all monthly observations are accumulated); and (2) the number of different species per event. As a measure of effort, I use the total number of different observers with at least two observations, in a given “event”. The final index is the average over all hexagons with at least one record, for a given year.

By changing the area of the hexagons, one is potentially changing results. This is the “modifiable area unit problem,” long-known to geographers (Openshaw 1984). Fortunately, in our case, the qualitative results are not affected by the resolution of the hexagons (correlations of data among resolutions are always above 0.7), and thus I shall report only on the analysis using the largest (two degrees) hexagons, of which there are 90.

The literature suggests that decrease in insect abundance is due to: (1) increased use of pesticides, (2) decreasing amount of habitat (or increasing amount of transformed land), and (3) climate change. At the scale of the whole country, I report regressions of data versus time series of pesticide use and deforestation rates.

## Methods

Citizen science data is not ideally suited to estimate trends, mostly because of the biased and uneven way sites, times and species are sampled (Isaac et al. 2014). In this work we try one of the three methods proposed by Isaac et al. (Isaac et al.): correcting the reported number of sights by a measure of effort.

iNaturalist data was downloaded from the Global Biodiversity Information Network (GBIF), as detailed below.

Data, then, was subsetted (keeping records with coordinates) for the four largest families in the Papilionoidea: Papilionidae (5,583 records), Pieridae (29,994 records), Nymphalidae (22,341 records) and Lycaenidae (2,326 records). The Lycaenidae was not used in the analysis due to the fact tha many species are of difficult to determine in the field. We also downloaded data from the genus Bombus (bumblebees, 6,543 records) and the two suborders of the Odonata (the Zygoptera, 12,772 records, and the Anisoptera, 24,003 records). For comparison purposes I also downloaded observations of the night shade family, the Solanaceae (39,014 records). The positions of every observation in Mexico are presented in the figures (S4 to S7) in the Supplementary Materials. I kept data tagged as “research quality” in the GBIF download, and performed a basic data-cleaning to keep coordinates inside Mexico. No attempt was made to correct for outdated taxonomy or other known issues present in aggregator’s data (Chapman 2005).

The data can be organized as a time-series, by pooling the observations in a year. Of course, this has the penalty of missing seasonality, but pooling by month produces tables that are too sparse, making them difficult to analyze. To assess the trends, including some measure of uncertainty, I took the averages of the calculated indices over all non-empty (I, e., with at least one observation) hexagons of two degrees of surface, and calculated its standard error.

Two indices are used: (1) different_species/observer and (2) observations/observer. The first is a measure of diversity, and the second a measure of abundance. I report on the two. Observers are the number of observers with at least two registered observations.

Since we wanted to summarize the trends, a useful statistic may be the slope of a linear model of index as a function of time, which call for regressions of index vs. year. However, as discussed before, the errors in many time-series problems are not independent, and the equal variance assumption of ordinary least squares is also often violated. If uncorrected, these problems “inflate” significance (McShane et al. 2019). Among many ways of dealing with the problem I choose performing generalized least squares regressions (Baillie and Kim 2018), which permits to include an autoregressive structure of correlations and violations of homoscedasticity. Two models were fitted to the data: an ordinary linear least squares, and a first order auto-regressive, moving average model (ARIMA) (Shumway and Stoffer 2005) allowing heteroskedasticity. The two models were compared using an ANOVA (Fox and Weisberg 2019) and the most likely one (based on the Akaike criterion), see Fox and Weisberg (2019) was used. This allows for a rigorous obtention of a probability for the observed values of slope under a null hypothesis of a slope equal to zero. Reporting “significance” of the slopes, is a practice that has been seriously criticized (McShane et al. 2019). Therefore, I simply report the probability value of the observation assuming a null model of no trend. If this probability is very small, I highlight the fact.

I included in the regressions two possible causal factors: forest loss and use of pesticides. The deforestation rate was obtained from the Global Forest Watch website (Sims et al. 2024) with a threshold of 30% of forest cover, as recommended by Sims et al. (2024). This dataset has maintained a methodological consistency (Hansen et al. 2013) that makes it preferable to the INEGI Series (Gebhardt et al. 2015). For use of agrochemicals I used the amount of pesticides used per hectare of cropland as reported in the FAO Web site. The data comes from government reports https://www.fao.org/faostat/en/#data/RP. For a discussion of the strengths and problems of the FAO dataset, see Shattuck et al. (2023).

Since in most cases the probability of the observed values of the slope of the index of diversity per unit of effort vs. time, under a null hypothesis of zero slope, was small, I assumed the regression could remove the time trend and looked for factors affecting just the residuals. In other words, I regressed the residual of the index vs. time regressions against two predictors: deforestation rate and use of pesticides. The results appear in the supplementary materials.

In order to aggregate by “biome” I choose the subdivision of the Potential Vegetation of Mexico (Rzedowsky 1986). A shapefile of Rezedowsky’s map, is available at CONABIO’s Geoportal, at scale 1:4,000,000, produced by Instituto de Geografía, UNAM México. I used this map to pool the iNaturalist records by Potential Vegetation, using the four categories with the highest number of iNaturalist reports.

An informal survey was circulated among scientists in three of the main Ecology research centers in Mexico (INECOL, Veracruz, Instituto de Ecologia, UNAM, and Ecosur, Chiapas). A total of 37 questionnaires were sent, using Qualtrics. The questions are reported in the Supplementary Materials section.

## Results

The informal questionnaire received 27 answers, out of 37 requests. Of the 27answers, 84% stated that they have noticed a decrease in the number of insects observed either in streetlights in the villages, or in the windshield or the radiator of the vehicles used to go to the field. Although without any statistical rigor, these answers suggest a widespread perception in field biologists in Mexico: insect populations are becoming smaller.

The iNaturalist data provides a more nuanced picture, but before looking at the behavior of biodiversity indices, it is good to have a look at some basic data. Indeed, both the number of species and the number of observers (with more than two observations) are growing, as shown in Figure (2)

**Figure (2).**
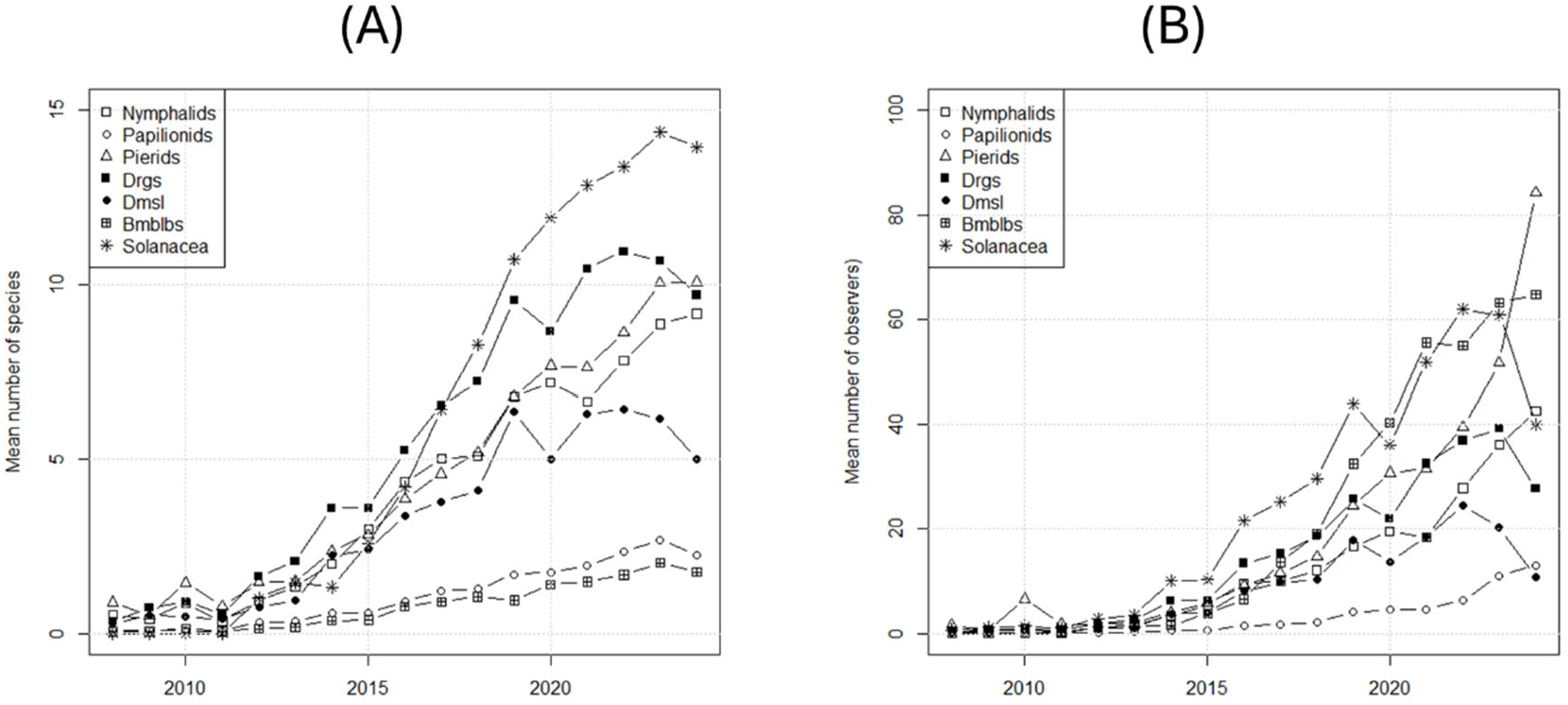
Growth in the mean number of species (A) and the mean number of observers with more than two observations, in the period. The average is taken over hexagons of two degrees of area covering the country. Drgs are the Anisoptera (dragonflies), and Dmsls are the Zygoptera (damselflies).

**Figure 2.**
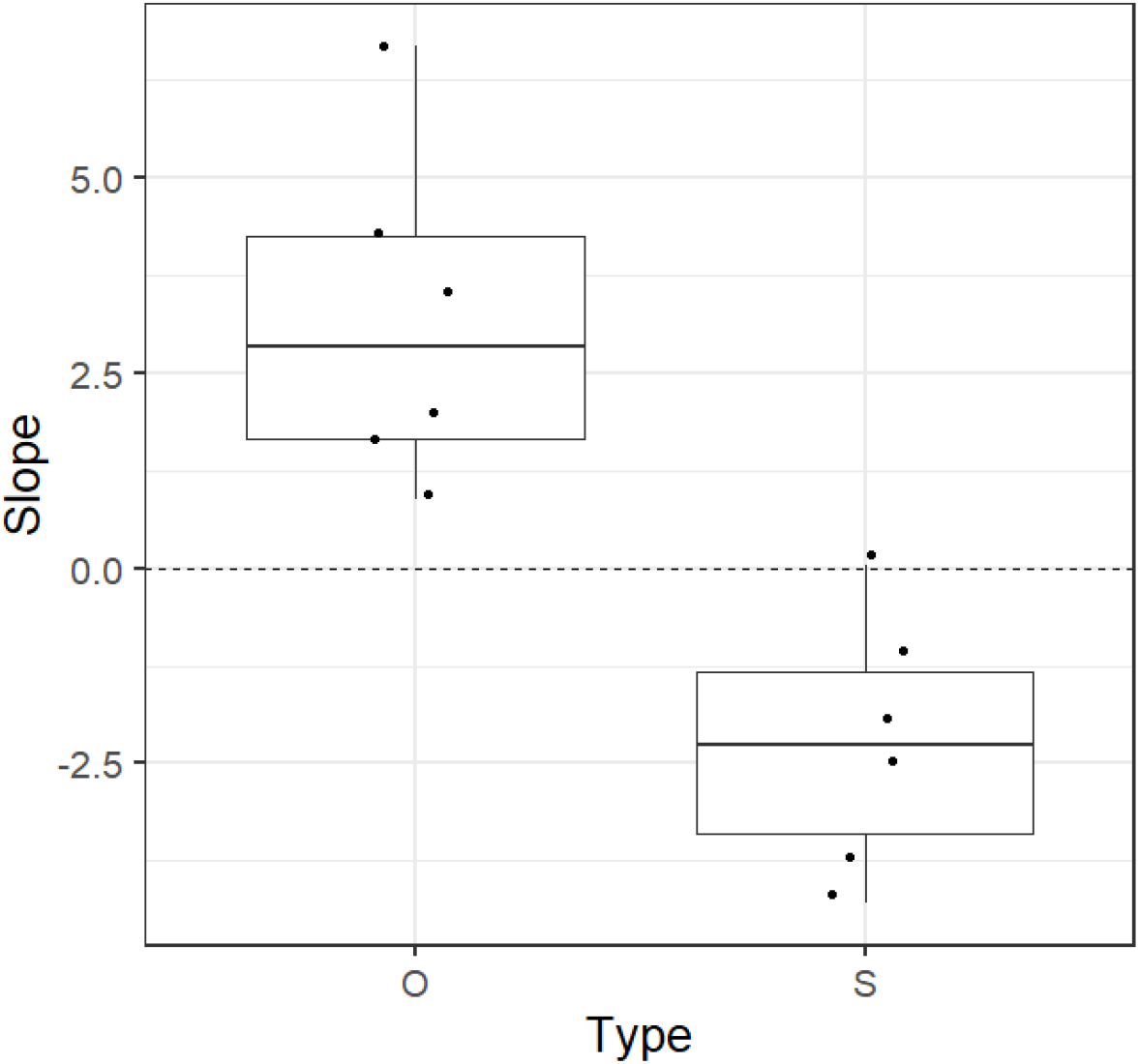
Box plot of the slopes of the regression of index vs. time, for Observations/effort (O) or Species/effort (S). Notice that for S five out of six slopes are negative. The dashed horizontal line highlights the zero slope.

Both observed species and observers grow in time. The fact that the number of observers grows introduces an important bias in the data, since it would be reasonable to expect more species (or more individuals) reported if there are more observers. However, although the **diversity** of insects appears to be diminishing, the evidence for a decrease in **abundance** is not clear. This is shown in figure (3) and tables (1) and (2) below.

**Figure 3.**
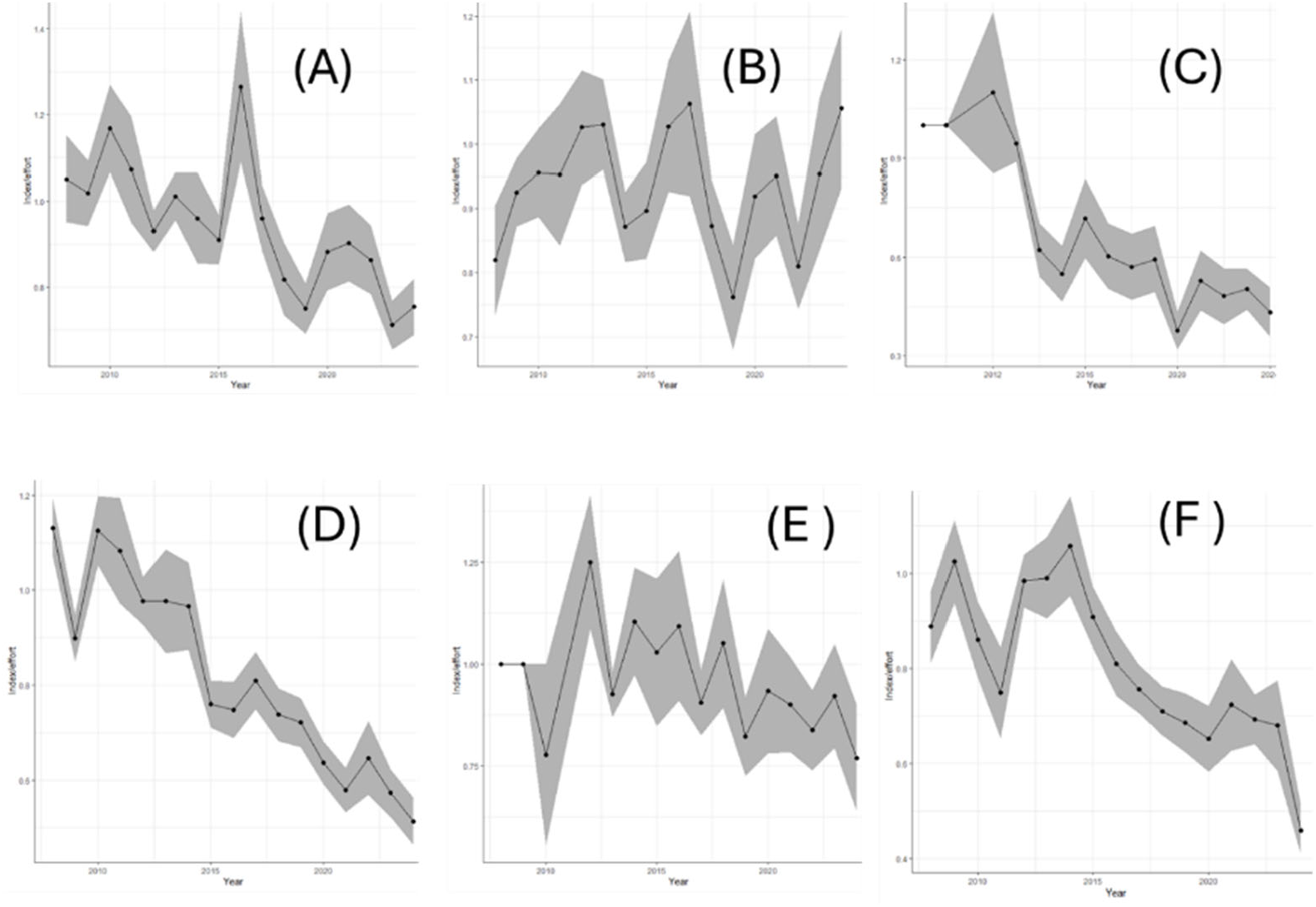
Average and standard error (band) of *different_species* per unit *effort* vs. time in years. for the bumblebees (A), the damselflies (Zygoptera) (B), the dragonflies (Anisoptera) (C), the nymphalids (D), the swallowtails (Papilionidae) (E) and the sulphurs (Pieridae) (F). With the exception of the damselflies, the slopes are all negative; and excepting (B) and (E), with very low probabilities of the observed values, under a null hypothesis of zero slope. This is reported in Table 2.

**Table 2.** Regression analysis (Generalized Least Squares) of diversity/observer vs. time for the iNaturalist data, for main taxonomic groups. The analysis was performed over the mean value in hexagons of 2 degrees of resolution. With the exception of the Zygoptera, where the first order autoregressive model did not converge, the ordinary least squares regression was not significantly different to models with autocovariance and heteroskedasticity and so the table shows the slope of ordinarly linear models of different_species/effort ∼ time. The probabilities of the obtained values under a null hypothesis of slope of zero are very small, with the exception of the dragonflies and the swallowtails (see figure 3)

| Species, 2 degrees |  |  |  |  |
| --- | --- | --- | --- | --- |
|  | Slope | p | Model | n |
| Bombus | -0.0428 | 0.0000237 | OLS | 6,543 |
| Anisoptera | -0.0206 | 0.00155 | OLS | 24,003 |
| Zygoptera | 0.0003 | 0.942 | OLS_NO_CNV | 12,772 |
| Nymphalidae | -0.0372 | 0.000000242 | OLS | 20,145 |
| Papilionidae | -0.0108 | 0.104 | OLS | 5,583 |
| Pieridae | -0.0247 | 0.000202 | OLS | 29,994 |

In table (2) I show that diversity per unit effort appears to decrease. However, the trends in the abundance (observations/number_observers) are either positive or indistinguishable from zero, as shown in Table 3.

In figure (2) we display boxplots of the slopes of the regressions for the two indices (species, and observations).

**Table 3.** Regression analysis (Generalized Least Squares) of records/observer vs. time for the iNaturalist data, for main taxonomic groups. The data was analyzed over hexagons of two degrees of resolution. Except for the Zygoptera, where a first order autoregressive model was used, the ordinary least squares regression was not significantly different to models with autocovariance and heteroskedasticity and so the table shows the slope of ordinary linear regressions of number_of_records/effort ∼ time. Notice that every regression has a positive, low probability slope.

|  | Slope | p | Model | n |
| --- | --- | --- | --- | --- |
| Bombus | 0.0446 | 0.00000766 | OLS | 6,543 |
| Anisoptera | 0.0201 | 0.00649 | OLS | 24,003 |
| Zygoptera | 0.0366 | 6.86E-08 | ARIMA | 12,772 |
| Nymphalidae | 0.0089 | 0.034 | OLS | 20,145 |
| Papilionidae | 0.0669 | 0.00000176 | OLS | 5,583 |
| Pieridae | 0.0154 | 0.0000984 | OLS | 29,994 |

The results above are very suggestive. It appears that diversity is decreasing, but abundance is stable. This is not in accordance with the informal perception of field biologists (as shown by the questionnaire), because most of them report an experience of diminished insect abundance. In the few cases in which the abundance of insects in Mexico are monitored systematically, one (the *Danaus plexippus*, the Monarch butterfly) is decreasing (Vidal and Rendón-Salinas 2014, Zylstra et al. 2021), whereas the *Anastrepha* fruit fly populations appear to be stable (Aluja et al. 2012, Ordano et al. 2013). Comparing these two cases is challenging, since the Monarch butterfly is affected by a variety of factors occurring at a continental scale, whereas the fruit flies perhaps are affected mostly by local factors.

Does the negative trend in diversity correlate with predictors often associated with insect loss? Forest cover as measured via remote sensing over a period of fifteen years (Hansen et al. 2013) is decreasing in Mexico (see Supplementary materials). Pesticide use per ha of crop, as reported by FAO, was increasing until 2018, when the FAO database shows an abrupt fall (see figure S8). The causes of the drop, if real, are not known, but after the COVID pandemic Mexico’s primary sector suffered a serious drop of activity (Sánchez et al. 2022). In any case, regressions of the residuals of the diversity/effort vs. time models against two predictors: deforestation rate and pesticide use per ha were never associated with small probabilities of an H0 of zero slope (see supplementary materials). This means that the data we used does not provide evidence that negative slopes in insect diversity are due to pesticide use or deforestation.

Finally, for the major taxonomic groups the slopes of the generalized least squares, in the first four potential vegetation types of Rzedowsky (1986) were most negative for Bumblebees in the Tropical Deciduous forest, followed by the Pine-Oak forest and then the Xerophitic shrub. For the butterflies, the most negative slope was in the pine-oak forest, followed by the tropical deciduous and the the Xerophitic Shrub (see Supplementary table S2).

## Discussion

The results I present of a diminishing trend of diversity but a stable pattern on abundance are compatible with several hypotheses. One, entirely biological, is that the diversity of insects is diminishing, although not the abundance. This suggests that the rarest species are disappearing, which means that the country is homogenizing (McKinney and Lockwood 1999). In other words, the very diverse and unique insect biodiversity of Mexico is slowly being replaced by a more homogeneous, more cosmopolitan set of species. This rather alarming hypothesis, supported by the citizen science data, needs to be more directly assessed in the field.

Another explanation for the negative trend observed in species numbers is that, with time, observers reach the asymptote of total number of species available to be observed. In other words, since the total number of species in any given area is roughly constant, with enough effort not more than that constant number can be reported, but if the number of observers is growing, one would see a negative trend in an index of species/observers. The total number of species in the database, for each taxonomic group, appears in table (1). In figure (1) we see that the average number of species reported per hexagon is well below that total. This suggests that the saturation effect is not present and the results presented here are indeed evidence of a diminishing trend in the diversity of insects. This complex point is discussed at more length in the Supplementary materials.

Finally, the negative trend would also be compatible with a hypothesis about the quality of iNaturalist observers: there are more observers, as the data shows, but they may change their discriminating capacity, or their interests, with time, perhaps because they focus on the common species. Unfortunately, the very nature of the information in citizen-science data makes it very difficult to assess this effect. This is in essence the major problem of unstructured citizen’s science data: methods are not standardized hence any trend in the data may be explained by a trend in the behavior of the observers.

What explanations can we have for no trends in the number of observations/effort? I can see two possibilities: one is that indeed more observers are reporting more individuals of common species. Another is that more observers simply means more observations and the ratio of observations to observers with more than 2 observations is roughly constant.

Having stated the above, it is always possible to accept the results I present as hypothesis to be inspected by more powerful methods. As such, the results very strongly suggest that the number of species in butterflies (important from a cultural perspective, and perhaps too as pollinators), the bumblebees (important as pollinators), and the Odonata (important as insect predators and as indicators) is decreasing. In other words, the biodiversity of Mexico is decreasing for some of the most important and underappreciated group of species. If confirmed, this would be an exceedingly alarming result. What may be the causes of this decrease? In study in Europe (Schuch et al. 2012), where a similar decrease in diversity was reported in a family of bugs of agricultural importance, the authors report a concomitant loss of non-agricultural habitat for the insects. These authors performed their study at the species level, and thus they can argue that the more specialized, less tolerant species are the ones disappearing due to the expansion of agriculture.

In view of the above, I regressed the residuals of least squares of the index vs. time against two factors related to agriculture: deforestation rate and use of pesticides, with data for the entire country. The results do not support a conclusion about the effect of deforestation or pesticides on the residuals (this approach removes the trend effect). However, it is known that species population dynamics is heavily affected by climate (Roy et al. 2001), and longer-term declines in insect abundance have been explained by climate, pesticides or changes in land-use (Basset and Lamarre 2019). The fact that I did not find significant effects of pesticides or deforestation in either one of the indices of biodiversity I used is likely due to a scale effect: the data on predictors is of national scale, whereas the observations in iNaturalist are point-like, although aggregated to 2 degrees. It would be necessary to obtain much higher resolution of the predictors to be able to meaningfully correlate insect indices with predictors.

Climate change is often cited as a cause of population decline in insects. However, it must be remembered that climate change is a long-term phenomenon that takes place at the scale of many decades. To demonstrate climate change as a factor affecting population size one needs to resort to modelling (Batalden et al. 2014) or to document the effect of mean and variance in climatic variables on long-term population time-series (Boggs 2016). The data that we use is not appropriate for this purpose.

Our results suggest that negative trends may not be the same in different ecological regions. However, it is very interesting that the iNaturalist data show that the Pine-Oak forest, the Xeric Shrub and the tropical deciduous forest may be the hotspots of loss of diversity. This may be a surprise given the almost universal concern about the tropical rain forests. Of course, another explanation would be the scarcity of data for the tropical wet vegetation types.

## Conclusion

Unstructured, un-systematic citizen’s data strongly suggest that insect diversity in Mexico is diminishing over time, at least for butterflies, bumblebees, and damsel and dragonflies, but insect abundance may be stable. However, the results are not unequivocal. Moreover, taken together they may be inconsistent with the informal field experience of many ecologists, and could be explained by assuming that observers change their behavior in time. Besides, evidence does not support that deforestation, which is increasing, nor use of pesticides, which until 2018 was also increasing, correlate with the loss of insect diversity.

Despite those caveats, the tens of thousands of observations of iNaturalist data for Mexico show a negative trend in insect diversity, for four conspicuous families of butterflies, for the Odonata, which are important insect predators(Simaika and Samways 2008) and for the bumblebees, a conspicuous and important group of pollinators (Stanley et al. 2015). Negative trends in insect abundance or diversity have been hypothesized to be caused by patterns of expansion of agriculture; widespread use of pesticides, and climate change. To prove that such factors are affecting insect abundance or diversity, more precise and long-term data would be required. Citizen’s science data provides a first glimpse of an alarming phenomenon, but it is not enough.

The inescapable conclusion is that Mexico needs to invest on insect monitoring schemes, countrywide, and based on systematic methodologies. This can be done using several approaches. The first, is to improve citizen’s science schemes, providing training and applying standard protocols, as it is done in Canada, the United States, and many countries of Europe (Streiter et al., 2024). This approach may only be useful for conspicuous, easily-identifiable species, but has a very substantial effect on the environmental awareness of the public (Dickinson et al. 2012), and if only for this reason, such efforts ought to be maintained.

There are also several methods resorting to advanced technologies, like computer vision, bioacoustics, and metagenomics (Van Klink et al. 2022). For bats, this is already being done in Mexico, by deploying automated acoustic sensors and then resorting to artificial intelligence to process the massive amount of data generated (Zamora-Gutierrez et al. 2020). Any of these methodologies require funding, training, and substantial analytical capacities.

Regardless of the method, Mexico, the fourth most biological diverse country in the planet, cannot afford not to monitor as many components of its biodiversity as possible.

## Data availability

The main data tables, and the R code are openly available (Creative Commons CC0: 1) in https://github.com/jsoberon/iNaturalistInsectsMexico

## Acknowledgements

I am very grateful to Luis Eguiarte and Rodrigo Medellin, of the Mexican *Instituto de Ecología* for their very acute and positive criticism of the methods I used. Their feedback made me think and change several times the way I analyzed the data. Exequiel Ezcurra, of the University of California at Irvine, and Carlos Martinez, formerly of the University of Wyoming, made very helpful statistical suggestions. My students Jennifer Ramos and Anahi Quezada helped me to download and organize data and patiently discussed the project with me. I am very grateful to those field ecologists in Mexico that took the time to answer my informal questionnaire about their experience with insects in the field.

## Notes

https://github.com/jsoberon/iNaturalistInsectsMexico

### Competing Interest Statement

The authors have declared no competing interest.

