## Supplementary materials for "Citizen science suggests decreased diversity of insects in Mexico, a megadiverse country"

SM1 Subdivision in hexagons

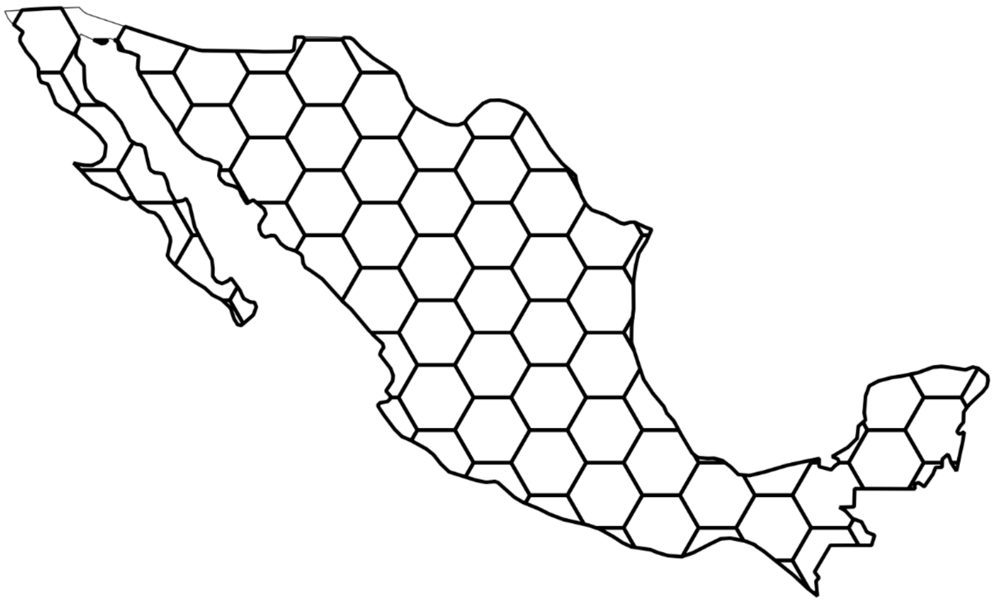

Figure S1. Hexagons of 2 degrees of resolution for Mexico. Obtained in Quantum GIS

SM2 Growth of iNaturalist

iNaturalist in Mexico is experiencing a substantial growth, that would bias the trends since every year there are more observers and observations, as shown in the table below

| | Butterflies | | Bombus | | Odonata | | Solanacea | |
|---|---|---|---|---|---|---|---|---|
| Year | Observers | Records | Observers | Records | Observers | Records | Observers | Records |
| 2008 | 41 | 186 | 2 | 3 | 20 | 58 | 0 | 0 |
| 2009 | 49 | 100 | 3 | 3 | 35 | 112 | 0 | 0 |
| 2010 | 67 | 605 | 9 | 9 | 48 | 124 | 0 | 0 |
| 2011 | 50 | 188 | 3 | 3 | 38 | 95 | 0 | 0 |
| 2012 | 104 | 276 | 8 | 10 | 66 | 246 | 39 | 104 |
| 2013 | 161 | 379 | 15 | 15 | 108 | 313 | 64 | 135 |
| 2014 | 221 | 735 | 30 | 48 | 151 | 852 | 65 | 133 |
| 2015 | 291 | 1049 | 37 | 69 | 199 | 903 | 97 | 327 |
| 2016 | 509 | 1869 | 76 | 163 | 328 | 1894 | 213 | 602 |
| 2017 | 657 | 2209 | 108 | 212 | 433 | 2217 | 376 | 1234 |
| 2018 | 811 | 2764 | 162 | 306 | 607 | 2670 | 763 | 1911 |

| | | | | | | | | |
|---|---|---|---|---|---|---|---|---|
| 2019 | 1222 | 4351 | 239 | 457 | 840 | 3981 | 1278 | 3347 |
| 2020 | 1401 | 5226 | 309 | 650 | 822 | 3332 | 1365 | 3966 |
| 2021 | 1577 | 5389 | 424 | 913 | 1075 | 4737 | 1993 | 5656 |
| 2022 | 2080 | 7214 | 566 | 1228 | 1200 | 5627 | 2250 | 5778 |
| 2023 | 2329 | 9395 | 517 | 1237 | 1271 | 5609 | 2541 | 6654 |
| 2024 | 2460 | 12950 | 501 | 1217 | 1279 | 4005 | 2713 | 7022 |

Table S1. Basic data for the four taxonomic groups for which indices of abundance were obtained.

### SM3 Theoretical value of the slope of diversity/observer vs. year

Since insects are increasingly reported in iNaturalist, we need to assess how much this increase may affect the index I use. In figure (2) we show a plot of the mean number of species reported, and observers, per year, in the period 2008 to 2024, for the butterflies, bumblebees and dragonflies and damselflies. The iNaturalist data show a simultaneous increase in the number of species observed and the number of observers. We are reporting species/observer. Given that the index is a quotient of two growing functions, what would be the theoretical value of its slope?

Assume that the observed number of species is a function of effort, with an asymptote, like in the Michaelis-Menten formula (Clench 1979) which is often used to model accumulation curves of species vs. effort. Thus, a model for the number of species observed, given the collecting effort, is s[f]=a*f/(1+b*f), where *f* is the effort, and *a* and *b* are parameters such that *a/b* is the total number of species in the region under consideration. The index I use is s[f]/f. Its derivative with respect to *f* is -*a***b*/(1+*b***f*)^2, which is always negative, regardless of the shape of the effort function. However, the precise value depends on the parameters a and b of the accumulation curves, and on the shape of the effort function. Since the total number of observed species, for large effort is *Tot = a/b*, the slope of the diversity/effort is -(a^2/(Tot*(1 + (a*f)/Tot)^2 )). This equation suggests that for large values of Tot the slope should be close to zero, and this is the assumption I make: the saturation effect should be small, except perhaps for the bumblebees. This problem deserves further consideration.

### SM4. Localities of iNaturalist reports

The localities of observations are illustrated in the following figures

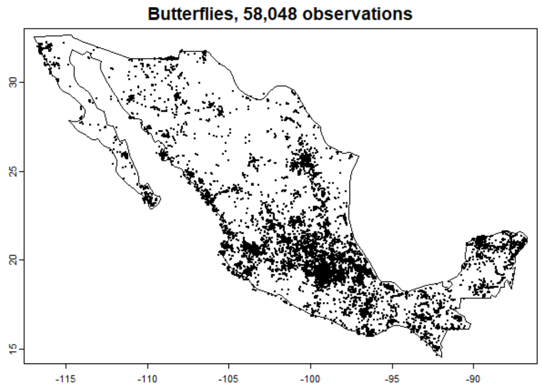


Figure S4. Localities in the map of iNaturalist for butterfly families (Nymphalidae, Papilionidae, Pieridae, Lycaenidae).

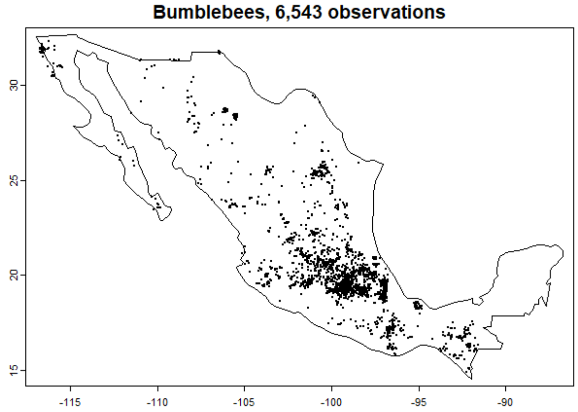


Figure S5. Localities in the map of iNaturalist for the genus Bombus.

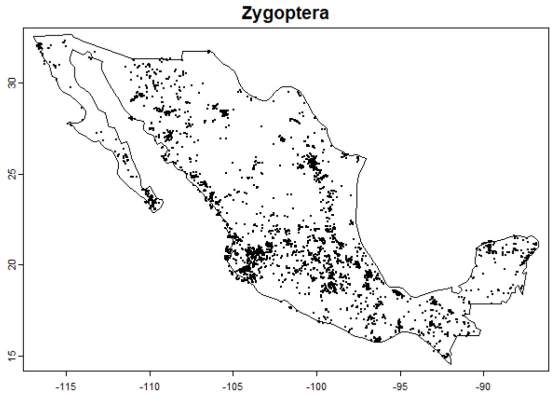


Figure S6. Localities in the map of iNaturalist for the sub order Zygoptera (damselflies).

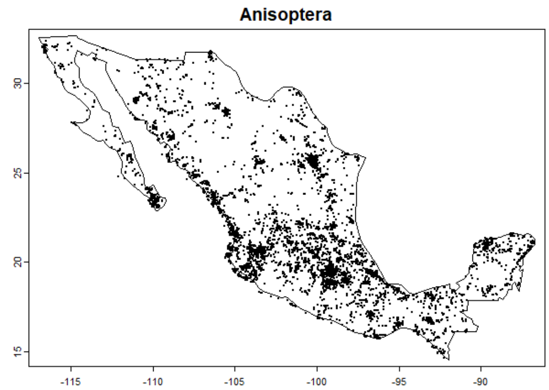


Figure S7. Localities in the map of iNaturalist for the sub order Anisoptera (dragonflies).

It is clear from the figures above that iNaturalist has a strong spatial bias, probably related to cities.

<u>SM5. Potential vegetation types of Rzedowsky</u>

The Potential Vegetation classes of Rzedowsky (1993) are:

Xerophitic shrubs (Mx)
Pine-Oak forest (Bce)
Grasslands and sabanas (P)
Tropical dry forest (Btc)
Tropical wet forest (Btp)
Spiny forest (Be)
Cloud forest (Bmm)
Tropical subdeciduous forest (Bts)
Mangrove and other wetlands (Vas)

And the percentage of pixels in the raster of Vegetacion Potencial de Mexico Rzedowsky, with iNaturalist observations are:

| | Mx | Bce | P | Btc | Btp | Be | Bmm | Bts | Vas | NumTot |
|---|---|---|---|---|---|---|---|---|---|---|
| VPR | 38.65 | 18.76 | 8.76 | 14.03 | 9.23 | 5.89 | 1.00 | 2.57 | 1.05 | 3,891 |
| Butterflies | 18.70 | 19.51 | 16.80 | 17.46 | 10.06 | 5.62 | 6.23 | 4.58 | 1.04 | 58,048 |
| Odonata | 27.44 | 14.37 | 15.31 | 23.10 | 6.28 | 4.80 | 2.09 | 6.16 | 0.46 | 36,775 |
| Solanaceae | 36.15 | 17.74 | 18.43 | 14.55 | 2.68 | 6.98 | 1.25 | 1.25 | 0.97 | 39,014 |
| Bombus | 17.94 | 33.23 | 30.54 | 4.02 | 2.78 | 3.90 | 5.12 | 0.11 | 1.50 | 6,543 |
| Mean | 27.78 | 20.72 | 17.97 | 14.63 | 6.21 | 5.44 | 3.14 | 2.93 | 1.00 | |

Table S2. Percentage of observations of different taxa in the Potential Vegetation map of Mexico. Ordered by mean proportion of observations. The first four vegetation types have at least 14% of the observations, but the last five types contain less than 7%. I used only the first four types. The first row is the percentage of area in the map, for that vegetation type.

It is clear that iNaturalist observations are not in proportion to the amount of vegetation type in the country. The grasslands and savannas have many more observations that what would be expected just by area. The regressions of diversity/effort vs. time in the four most visited vegetation types appear below.

| Taxon | Potential Vegetation | Slope X 1000 | p | p(ARIMA) | n | Model |
|---|---|---|---|---|---|---|
| Bombus | All | -10.31 | 2.700E-03 | 1.040E-03 | 6,543 | ARIMA |
| Bombus | MX | -16.94 | 9.000E-03 | 6.400E-05 | 1,174 | ARIMA |
| Bombus | POF | -18.28 | 6.440E-10 | 1.110E-06 | 2,174 | ARIMA |
| Bombus | G | -2.52 | 6.800E-01 | 4.100E-01 | 1,998 | OLS |
| Bombus | TDF | -21.39 | 1.600E-02 | 8.760E-01 | 263 | OLS |
| Butterflies | All | -6.11 | 3.123E-03 | 2.740E-02 | 58,048 | ARIMA |
| Butterflies | MX | -8.72 | 3.510E-02 | 2.550E-02 | 10,857 | ARIMA |
| Butterflies | POF | -10.17 | 1.530E-01 | 5.420E-03 | 11,327 | ARIMA |
| Butterflies | G | -5.02 | 3.280E-01 | NoConvergence | 9,750 | |
| Butterflies | TDF | -9.34 | 1.150E-02 | 5.400E-02 | 10,133 | ARIMA |
| Odonata | All | -0.89 | 6.800E-01 | 5.400E-02 | 36,775 | ARIMA |
| Odonata | MX | 8.06 | 3.760E-02 | NoConvergence | 10,090 | |
| Odonata | POF | -6.62 | 5.120E-02 | 9.010E-01 | 5,283 | OLS |
| Odonata | G | 1.44 | 8.430E-01 | 4.200E-02 | 5,630 | ARIMA |
| Odonata | TDF | -7.85 | 4.200E-02 | 1.430E-01 | 8,495 | OLS |
| Solanaceae | All | 3.13 | 1.650E-01 | 7.930E-01 | 39,014 | OLS |
| Solanaceae | MX | 1.97 | 4.440E-01 | 7.810E-01 | 14,105 | OLS |

| | | | | | | |
|---|---|---|---|---|---|---|
| Solanaceae | POF | 1.1 | 6.820E-01 | 7.810E-01 | 6,920 | OLS |
| Solanaceae | G | -16.25 | 2.000E-02 | 8.880E-01 | 7,189 | OLS |
| Solanaceae | TDF | 8.71 | 1.040E-01 | 8.880E-01 | 5,676 | OLS |

Table S3. Results of GLS regressions of different_species/unit_effort for the four most common Potential Vegetation types, aggregated by two degrees hexagons. Slope X 1000. The number of observations for the taxon in the Potential Vegetation class is *n*.

In the table below I report the number of species

| Taxon | VPR | Total Names | Reported Names |
|---|---|---|---|
| Bombus | Mx | 16 | 10 |
| Bombus | POF | 20 | 15 |
| Bombus | G | 9 | 8 |
| Bombus | TDF | 14 | 7 |
| Butterflies | Mx | 153 | 44 |
| Butterflies | POF | 220 | 40 |
| Butterflies | G | 86 | 24 |
| Butterflies | TDF | 198 | 32 |
| Odonata | Mx | 185 | 36 |
| Odonata | POF | 187 | 60 |
| Odonata | G | 105 | 56 |
| Odonata | TDF | 184 | 65 |
| Solanaceae | Mx | 176 | 46 |
| Solanaceae | POF | 210 | 36 |
| Solanaceae | G | 111 | 24 |
| Solanaceae | TDF | 153 | 36 |

Table S4. Number of species in the iNaturalist database, for different taxa in the four most visited potential vegetation types of Rzedowsky (third column) and maximum number of species reported by an observer (with more than two observations) for each combination of taxon and vegetation type. With the exception of Bombus, every taxon is under sampled in the vegetation types used.

SM6. Regressions over the entire country

Below I show graphs of the indices of number of species/effort for four families of butterflies.

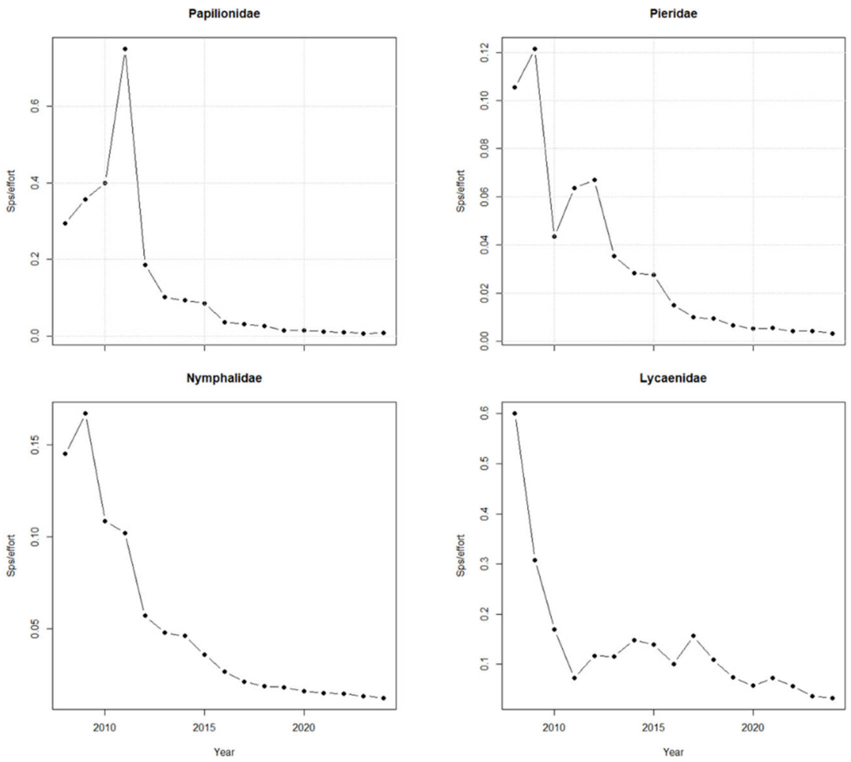


Figure S5. Species/observer vs. year for four families of butterflies. Data aggregated for all Mexico. In all cases the slope is negative.

| | Slope | p | n | Model used |
|---|---|---|---|---|
| Papilionidae | -2.63 | 0.282 | 5538 | ARIMA |
| Pieridae | -6.26 | 0.000000088 | 29994 | OLS |
| Nymphalidae | -1.0 | 0 | 20145 | ARIMA |
| Lycaenidae | -10.0 | 0.0025 | 2316 | ARIMA |

Table S6. Generalized least squares regressions for the species/effort vs. year for the four butterfly families, aggregated over all Mexico. The slopes are multiplied by 1,000. For three of the four families, performing a regression with a first-order autocorrelated structure was better than performing an ordinary least squares.

SM7. Predictors of loss of diversity, and results of the regressions

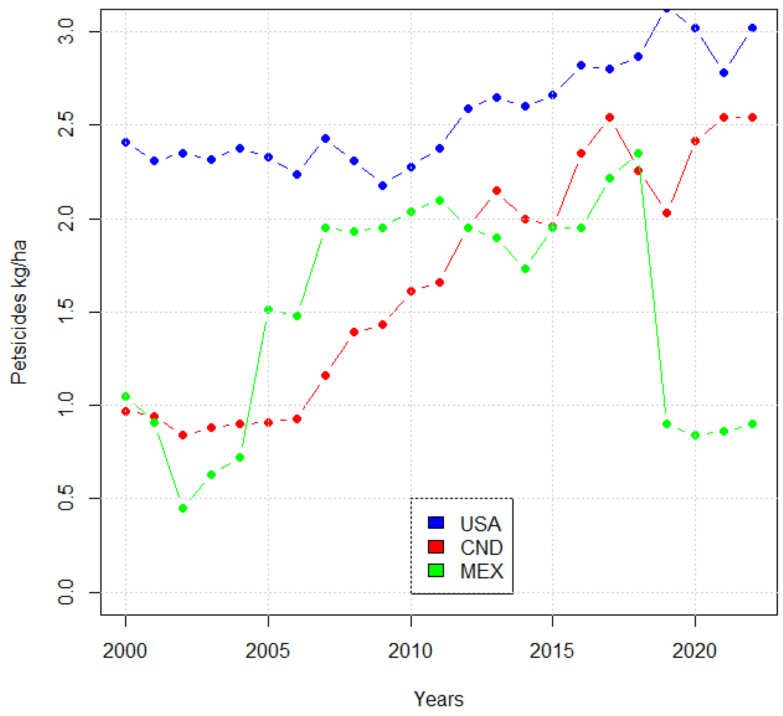


Figure S8. Rates of pesticide use (kg/ha) in the three largest North American Countries, as reported in the FAO world statistics database (https://www.fao.org/statistics/en ). USA is the United States, CND is Canada, and MEX is Mexico.

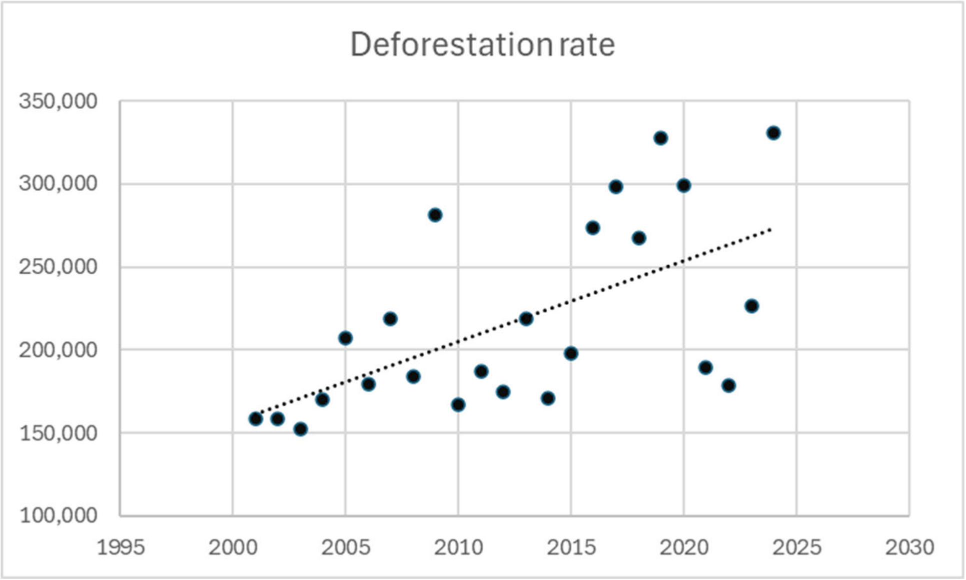


Figure S9. Loss of forest cover, for Mexico, in ha (https://www.globalforestwatch.org/ ).

Although for all the insect groups the regressions on the residuals of the GLS (of index vs. year) against deforestation and pesticide use were negative, they never had a very low probability under the null hypothesis. This means that the data does not provide evidence of a relationship between the negative trend in diversity and the predictors. Table S3 shows the results for hexagons of two degrees.

| Butterflies | Value | p-value |
|---|---|---|
| (Intercept) | 0.0166 | 0.588 |
| DefRate | -0.0456 | 0.161 |
| Pesticide Use | -0.0542 | 0.129 |

| Bombus | Value | p-value |
|---|---|---|
| (Intercept) | 0.018 | 0.848 |
| DefRate | -0.0227 | 0.814 |
| Pesticide Use | -0.0113 | 0.914 |

| Odonata | Value | p-value |
|---|---|---|
| (Intercept) | -0.00538 | 0.743 |
| DefRate | -0.01159 | 0.493 |
| Pesticide Use | -0.0036 | 0.843 |

| Solanaceae | Value | p-value |
|---|---|---|
| (Intercept) | -0.0444 | 0.473 |
| DefRate | 0.0157 | 0.774 |
| Pesticide Use | 0.1131 | 0.183 |

Table S7. Regressions of residuals vs. standardized predictors (Deforestation rate: DefRate) and Pesticide use per hectare. No slope has low probability under an H0 of zero slope.

<u>SM8. Informal questionnaire</u>

The questionnaire had the following results:

| **Do you do field work?** | Number of Answers |
|---|---|
| Less than annually | 1 |
| Annually | 12 |
| Monthly | 13 |

| **How long ago did you start going to the field** | |
|---|---|
| Five or fewer years | 0 |
| Ten or less years | 1 |
| More than ten years | 25 |

| **Have you noticed changes in insect abundance** | |
|---|---|
| No | 2 |
| Yes, an increase | 1 |
| Yes, a decrease | 23 |

| **How have you noticed?** | |
|---|---|
| Insects impacted on vehicules | 19 |
| Clouds of insects aronund lights | 17 |
| Direct monitoring | 10 |
| Other | 10 |

| **How much you do trust your observations?** | |
|---|---|
| Very reliable | 14 |
| Reliable | 6 |
| Doubtful | 6 |

There was a total of 27 answers, out of 37 requests. Notice that the questions are not exclusive, so the sum of the answers may not be 27.
